# Identification, structure-activity relationship and *in silico* molecular docking analyses of five novel angiotensin I-converting enzyme (ACE)-inhibitory peptides from stone fish hydrolysates

**DOI:** 10.1101/317198

**Authors:** Shehu Muhammad Auwal, Najib Zainal Abidin, Mohammad Zarei, Chin Ping Tan, Nazamid Saari

**Affiliations:** Department of Food Science, Faculty of Food Science and Technology, Universiti Putra Malaysia, 43400 Serdang, Selangor, Malaysia; (S.M.A.), (N.Z.A), (M.Z.); Department of Food Technology, Faculty of Food Science and Technology, Universiti Putra Malaysia, 43400 Serdang, Selangor, Malaysia; (T.C.P.); Department of Biochemistry, Faculty of Basic Medical Sciences, Bayero University, Kano, Nigeria; Department of Food Science and Technology, College of Agriculture and Natural Resources, Sanandaj Branch, Islamic Azad University, Sanandaj, Iran

**Keywords:** ACE-inhibitory hydrolysates, stone fish, peptides, hydrophobicity, isoelectric properties, molecular docking

## Abstract

Stone fish is an under-utilized sea cucumber with many health benefits. Hydrolysates with strong ACE-inhibitory effects were generated from stone fish protein under the optimum conditions of hydrolysis using bromelain and fractionated based on hydrophobicity and isoelectric properties of the constituent peptides. Five novel peptide sequences with molecular weight (mw) < 1000 daltons (Da) were identified using LC-MS/MS. The peptides including ALGPQFY (794.44 Da), KVPPKA (638.88 Da), LAPPTM (628.85 Da), EVLIQ (600.77 Da) and EHPVL (593.74 Da) were evaluated for ACE-inhibitory activity and showed IC_50_ values of 0.012 mM, 0.980 mM, 1.31 mM, 1.44 mM and 1.68 mM, respectively. The ACE-inhibitory effects of the peptides were further verified using molecular docking study. The docking results demonstrated that the peptides exhibit their effect mainly via hydrogen and electrostatic bond interactions with ACE. These findings provide evidence about stone fish as a valuable source of raw materials for the manufacture of antihypertensive peptides that can be incorporated to enhance therapeutic relevance and commercial significance of formulated functional foods.

## Introduction

Public adherences to unhealthy diet and poor lifestyle have been inflicted among the risk factors associated with etiology of hypertension. Consequently, increased awareness about the significance of healthy diet has led to the recent development of biologically active peptides through enzymatic hydrolysis of food proteins. The peptides consists of specific amino acids sequences and perform at least one or more physiological role on the body system including; antihypertensive activity (1 - 4), antioxidant activity (5 - 6), antibacterial activity (7), anti-inflammatory and opioids activities (8).

Angiotensin I-converting enzyme plays a primary role in the rennin angiotensin and kinin-kallikrein systems by increasing the production of angiotensin II and decreasing the formation of bradykinin thereby resulting in hypertension. The food derived antihypertensive peptides have been associated with safe effects and exhibit their role through the inhibition of ACE (9, 10). Thus, food peptides can be used as alternative to synthetic ACE-inhibitors such as enalapril and lisinopril which are often accompanied with certain adverse effects e.g thirst, diarrhea, skin rashes, fever and headache (11, 12). Many potent ACE-inhibitory peptides have been previously generated through protease-aided hydrolysis of food proteins obtained from both animal and plant sources including those of marine invertebrates (12 - 15). The protease-generated ACE-inhibitory peptides are in form of mixture (protein hydrolysates) that needs to be purified for the identification and structure-activity relationship study of the potent sequences.

A series of chromatographic techniques including reverse phase-high performance liquid chromatography (RP-HPLC), isoelectric focusing (IEF)-electrophoresis and QTOF-LC/MS mass spectrometry are used in the fractionation of complex mixture of peptides to ease identification of the potent sequences (4, 16).

The structure-activity relationship is studied to deduce the combination of COOH and NH_2_ terminal tripeptides of the identified sequences which are responsible for their observed inhibitory effect on ACE. Moreover, the study for the nature of interaction between these peptides sequences and catalytic site of ACE is accomplished by molecular docking study to unravel more of their desirable properties for therapeutic and commercial applications. The under-utilized stone fish used in this study is a marine invertebrate belonging to the sea cucumber species that is rich in biologically active compounds (17). Hydrolysates with strong ACE-inhibitory effects have been generated when stone fish protein was hydrolyzed using bromelain under the optimum conditions of pH, temperature, enzyme/substrate ratio (E/S) and time (2). However, the hydrolysates constitute the whole mixture of peptides that needs to be purified for the identification of the potent ACE-inhibitory sequences. Hence, we fractionated the ACE-inhibitory hydrolysates produced under the optimum hydrolysis conditions and identified the potent peptides present. We further elucidated the structure activity relationship and molecular mechanism through which the peptides exhibit their inhibitory effects on ACE.

## Materials and Methods

### Materials

The stone fish was collected from langkawi breeding center in Malaysia during low tide. Internal organs were removed and the tissue was washed before packaging in plastic containers. The packaged tissue was taken to the lab and rewashed prior to their storage at - 80°C. The frozen tissue was then lyophilized, grinded and sieved through a 600 μm wire mesh followed by storage at −40°C before hydrolysis.

Commercial food grade bromelain (from Pineapple stem, 2.4 to 3 U/mg) was purchased from Acros Organics (Geel, Belgium). Potassium salts (mono and dibasic) were obtained from Merck KGaA (Darmstadt, Germany). N-hippuryl-Histidyl-Leucine tetrahydrate (HHL) and angiotensin I-converting enzyme (from rabbit lung) were purchased from Sigma aldritch (St. Louis, Mo, USA). All other chemicals and reagents used were of analytical grade and supplied from Fisher Scientific (Loughborough, Leics, UK) and Merck KGaA (Darmstadt, Germany).

### Preparation of Stone fish protein hydrolysates (SFPH)

The ACE-inhibitory hydrolysates were produced under the optimum conditions for the hydrolysis of stone fish protein with bromelain (pH 7, Temp 40oC, E/S 2% and time 240 min) according to Auwal et al. (2). The substrate was mixed with the enzyme in a glass flask with the phosphate buffer (50mM, pH 7) and incubated in a water bath shaker at 150 rpm under the optimum temperature condition. After 240 min, the mixture was boiled at 100°C for 10 min to in activate the enzyme and the peptide mixture (hydrolysates, mwt <10,000 Da) were separated by centrifugation at 10, 000 × g, at 4°C for 20 min), lyophilized and stored at −80°C before used.

### Hydrophobicity-based fractionation using RP-HPLC

The lyophilized sample of Bromelain-generated stone fish protein hydrolysates was prepared at a concentration of 100 mg/mL with mobile phase A (made up of 0.1% trifluoroacetic acid (TFA)) in deionized water. The solution was then filtered through a 0.2 μm nylon membrane and loaded at 500 μL into a semi preparative RP-HPLC column (C18, 9.4 × 250 mm, Technologies, Santa Clara, USA), that was previously conditioned with mobile phase A. The peptides were eluted from the hydrolysates or their mixture at a flow rate of 4 mL/min by mobile phase A and mobile phase B into a total of 45 fractions of 6 mL each. The mobile phase B contained 0.1% TFA in acetonitrile (ACN). The fractions were collected within a total period of 67.5 min with only mobile phase A being used during the first 10 min and only mobile phase B during the last 12.5 min. The absorbance of each eluted fraction was monitored at a wavelength of 205 nm. Each fraction was lyophilized, reconstituted in 300 μm borate buffer (50 mM, pH 8.3) and assayed for ACE-inhibition.

### Isoelectric properties based fractionation

The RP-HPLC fractions with the highest ACE-inhibitory effects were selected and separated further based on the isoelectric points of their constituent peptides by IEF-electrophoresis using Agilent 3100 OFFGEL fractionator (Agilent Technologies, Santa Clara, USA). The pH graded gel strips were initially rehydrated and fixed on to the separating plate. Then, 150 μL of the sample was transferred to each well and the separation was achieved at high voltage (500–4000 V). At the end of the process, the content of each well was then gently withdrawn and stored at −80°C for subsequent analysis.

### Peptide Sequencing Protocol Using Q-TOF LC/MS

The tubes containing the IEF sub-fractions with the highest ACE-inhibition were high vortexed and spun at 15 000 rpm for 10 min. The supernatant was then separated and analyzed for peptides, using an ultra-high performance liquid chromatography system (Agilent 1200 HPLC-Chip/MS Interface, Santa Clara, USA) coupled to a high-resolution, accurate mass hybrid quadrupole-time of flight (Q-TOF) mass spectrometer (Agilent 6550 iFunnel Q-TOF LC/MS, Santa Clara, USA). In brief, a 1 μL from each peptide sample was injected in to an Agilent Large Capacity Chip, 300 Å, C18, 160nL enrichment column & 75umx150mm analytical column (P/N: G4240-62010) and separated against a gradient elution of ACN in 0.1% formic acid within 5% to 75% for 39 min. The peptides were monitored in an MS-mode within the range of 110 and 3000 m/z and selected automatically for collisionally induced dissociation (CID, MS/MS) that was based on charge state preference: 2, 3, > 3.

### Database Searching

The available data obtained from the samples by the Q-TOF-MS were searched against the animal species sub-directory of the SwissProt.Holothuridae.January2017.1047.fasta database (UniProt, EBI, UK) using the SpectrumMill search engine (SpectrumMill Rev.B.04.00.127, Agilent Technologies). Basic parameters were set at a precursor mass tolerance of 10 ppm, a product mass tolerance of 50 ppm and a ‘no-enzyme’ constraint. Peptides with score above 6 and a percentage of scored peak intensity higher than 50% were considered a match.

### ACE-inhibitory activity assay and IC_50_ determination

The peptides were chemically synthesized through solid phase synthesis for peptides ≥5amino acids with a percentage purity of > 90% and supplied by Genscript (New Jersey, USA). The ACE-inhibitory activity was measured according Jimsheena & Gowda (18) with minor modifications. The Peptides stock solutions were diluted over ranges of concentration for each peptide. A 15 μL of the peptide was pre-incubated with 10 μL of ACE (100 mU/mL) at 37°C for 10 min. Then 50 μL of the 5mM HHL substrate containing 0.3 M NaCl in 50 mM borate buffer pH 8.3 was added and the incubation was continued for the next 60 min. The reaction was stopped using 1 M HCl. Benzene sulfonyl chloride (BSC) and pyridine were then added at 75 μL and 150μL, respectively and the amount of hippuric acid produced was quantified by the intensity of the yellow color produced in a microplate reader at 410nm. The ACE-inhibitory activity of the peptides was then calculated from the average of three values as follows:

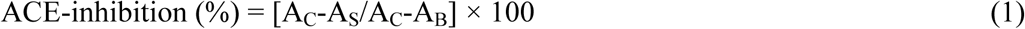

Where A_C_ means absorbance of control (where ACE and substrate were present), A_S_ means absorbance of sample (where peptide, ACE and substrate were present), while A_B_ means the absorbance of blank (where only substrate was present).

The IC_50_ value was defined as the concentration of peptides (inhibitor) required to exhibit 50% of ACE-inhibitory effect and was calculated from a non-linear regression plot of ACE inhibition (%) against peptide concentration (mM) using Graphpad Prism 7 software (GraphPad Software Inc., California, USA).

### *In silico* molecular docking of the inhibitory peptides and ACE

The preliminary step involved preparation of ACE as the protein molecule and peptides as the ligands so as to minimize docking errors due to incorrect assignments.

A Protein Preparation Wizard (Maestro, version 10.2, Schrödinger, LLC, New York, USA) was used to fully prepare an all atom protein model of the ACE from an X-ray crystallographic structure of human-testicular ACE-captopril complex (PDB ID: 1uzf) obtained from the Research Collaboratory for Structural Bioinformatics Protein Data Bank (RCSB PDB, http://www.rcsb.org/pdb/home/home.do). Then, the Prime software (Prime, version 4.0, Schrödinger, LLC, New York, USA) was used for loops and missing side chains filling. Atomic molecular valency was attained through hydrogen addition and the optimal hydrogen bond network was achieved according to the following steps; (a) The hydroxyl and thiol groups were rearranged as their orientation couldn’t be predicted from the x-ray protein structure (b) The terminal amide groups in asparagine (Asn) and glutamine (Gln) and an imidazole ring in histidine (His) were flipped at 180° to improve their charge-charge interaction with neighboring groups (c) The protonation and tautomeric state of His atom were predicted based on Schrödinger Software Team, 2015. Protein minimization and water molecules removal was carried out in order to relieve stearic effect. Then an Impact Refinement module (Impref) (Impact, version 6.7, Schrödinger, LLC, New York, USA) was used to restrain all heavy atoms to a minimum degree of deviation of the final result to the input geometry. The latter was set at maximum energy convergence, or the root-mean-square deviation cut-off value of 0.30 Å and force field of OPLS-2005 based on an optimized potentials for liquid simulation.

Captopril and the five potent stone fish protein hydrolysates-derived peptides were selected as ligands. All the peptides were managed using energy minimizing module while the captopril was prepared by LigPrep 3.4 version (Schrödinger, LLC, New York, USA). Although for each ligand there were maximum number of 32 stereoisomers and tautomers being generated, only one ring conformation of low energy was selected per ligand to accomplish the docking analysis.

A GLIDE extra precision software version 6.7 (GlideXP, Schrödinger, LLC, New York, NY, 2015) was employed to perform the docking in which docking parameters were maintained as default with no bonding constraints assign in the course of calculations. An empirical E-scoring function which approximates the ligand binding free energy was used to predict the potential energies of the docked molecules while post docking was performed with OPLS 2005 force field and one pose was saved per ligand. The binding affinity between the ACE and peptides were determined as negative values of GLIDE scores (kj/mol), such that the higher the negative value of the pose the stronger the protein-ligand (ACE-peptide) interaction. The docked poses obtained were further analyzed by GLIDE viewer mode and compared among the ligands with respect to their GLIDE docking score and binding affinity.

### Statistical analysis

Statistical analysis was carried-out using the Minitab (16.0 software version) from Minitab Inc. (State College, Pennsylvania, USA). The data are summarized as mean±standard deviations from triplicate observations. Significant difference was then determined using Tukey’s significance test, at p < 0.05.

## Results and Discussion

### Fractionation of Stone Fish Protein Hydrolysates Produced under Optimum Hydrolysis Conditions and Analysis of Peptide Sequence and ACE-inhibitory Activity

The stone fish-derived ACE-inhibitory hydrolysates generated under the optimum conditions of pH 7, temperature 40°C, E/S 2% and time 240 min were sequentially fractionated based on their hydrophobicity and isoelectric properties using RP-HPLC and IEF-electrophoresis, respectively. The constituent peptides were then identified by QTOF-LC/MS and evaluated for ACE-inhibitory effects.

Firstly, a semi-preparative reverse-phase high performance liquid chromatography (RP-HPLC) was used to separate the peptides mixture based on their hydrophobicity. The selection of this technique was because of its reproducibility due to sensitivity and high resolution (19, 20). The peptides were separated on a C_18_ column into 45 fractions using a system of semi-preparative RP-HPLC. Peptides separation was accomplished by gradient elution against mobile phase A containing 0.1% TFA in deionized water and mobile phase B consisting of 0.1% TFA in ACN at a flow rate of 4 mL/min. The concentration of mobile phase B was 0.0% during the first 10 min. It then increased linearly to reach 100% at 67.5 min (Figure 1a). More hydrophilic peptides fractions were collected at the early stage while highly hydrophobic peptides fractions were separated at later stage. The ACE-inhibitory activity of each fraction was determined as previously described.

**Figure 1.**
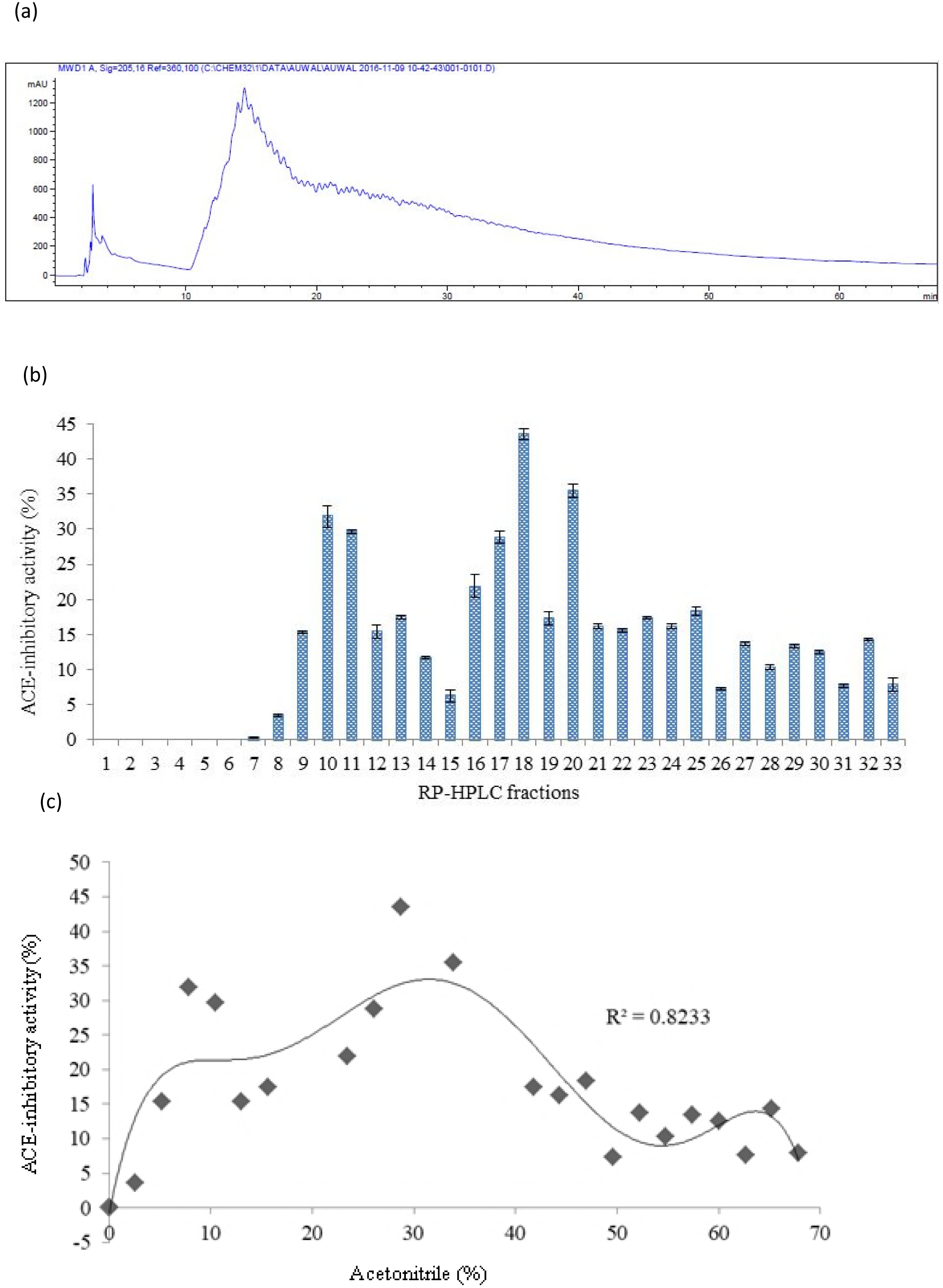
Hydrophobicity-based fractionation of bromelain-generated stone fish protein hydrolysate (a) Chromatogram of semi-preparative RP-HPLC; (b) ACE-inhibitory activities of each of the potent fraction; (c) Relation between ACE-inhibitory activity and percentage acetonitrile being used.

As shown in Figure 1b, no ACE-inhibitory activity was detected in the first 7 fractions. The first ACE-inhibitory activity of 3.55% was observed in fraction 8 which was eluted at 2.61% ACN gradient. The fractions 10, 11, 17, 18 and 20 exhibited relatively highest ACE inhibitory activity of 31.83%, 29.67%, 28.81%, 43.50% and 35.45% and were eluted against ACN gradient of 7.83%, 10.44%, 26.10%, 28.71% and 33.93%, respectively (Figure 1c). The last ACE-inhibitory activity of 7.86% was detected in fraction 33 eluted against 67.86% ACN gradient. Thereafter no ACE-inhibitory potency was observed with increase in hydrophobicity and elution time. Hence, hydrophobicity plays a significant role on the ACE-inhibitory effect of the peptides. These findings are in agreement with the previous observation reported by Yea et al. (16) where hydrophobicity was shown to affect the ACE-inhibitory activity of peptides. Moderately hydrophobic peptides were most potent and demonstrated highest ACE-inhibitory effect. Highly hydrophilic and highly hydrophobic peptides revealed no inhibitory effect against ACE. Accordingly, high hydrophilicity of peptides was reported to halt their interaction with active site of ACE resulting in low or no inhibitory effect on the ACE (21, 22). Consequently, the absence of ACE-inhibitory effect in the peptide fractions 1-7 might be related to high content of hydrophilic amino acids in their sequences. Similarly, fractions eluted near 100% hydrophobicity might constitute a mixture of unhydrolyzed proteins with no peptide being released to exhibit inhibitory effect against ACE (16).

Secondly, the RP-HPLC fractions 10, 11, 17, 18, and 20 with the highest ACE-inhibitory activity were selected and further fractionated based on their isoelectric properties by a system of OFFGEL fractionation using IEF-electrophoresis (Table 1).

**Table 1.** ACE-inhibitory effect of the RP-HPLC fractions and IEF sub-fractions

| RP-HPLC fractions | ACE-inhibitory activity (%) | IEF sub-fractions | ACE-inhibitory activity (%) |
| --- | --- | --- | --- |
| 10 | 31.83±1.570<sup>c</sup> | 11 | 64.49±0.417<sup>b</sup> |
| 11 | 29.67±0.290<sup>d</sup> | 9 | 69.21±0.325<sup>a</sup> |
| 17 | 28.81±0.850<sup>d</sup> | 1 | 43.92±0.134<sup>e</sup> |
| 18 | 43.50±0.778<sup>a</sup> | 2 | 60.43±0.778<sup>c</sup> |
| 20 | 35.45±0.919<sup>b</sup> | 9 | 52.11±1.131<sup>d</sup> |

Results are expressed as mean ± SD of triplicate determinations. Means of ACE-inhibitory activity (%) which carry different superscript letters are significantly different at $P \leq 0.05$ .

Peptides with similar isoelectric point (PI) were separated as clusters on an immobilized gel strip along a gradient of pH 3–10. Separated peptides with PI in the acidic (pH 3.6 - 6.5) and basic compartments (pH 7.7 – 10.0) showed higher ACE-inhibitory activity compared to those separated at neutral point (Figure 2). This showed that the charge nature of the acidic and basic peptides might have contributed to their observed ACE-inhibitory activity since charge distribution affects complementarity and binding of an inhibitor to the active site of an enzyme to form an enzyme– inhibitor complex.

**Figure 2.**
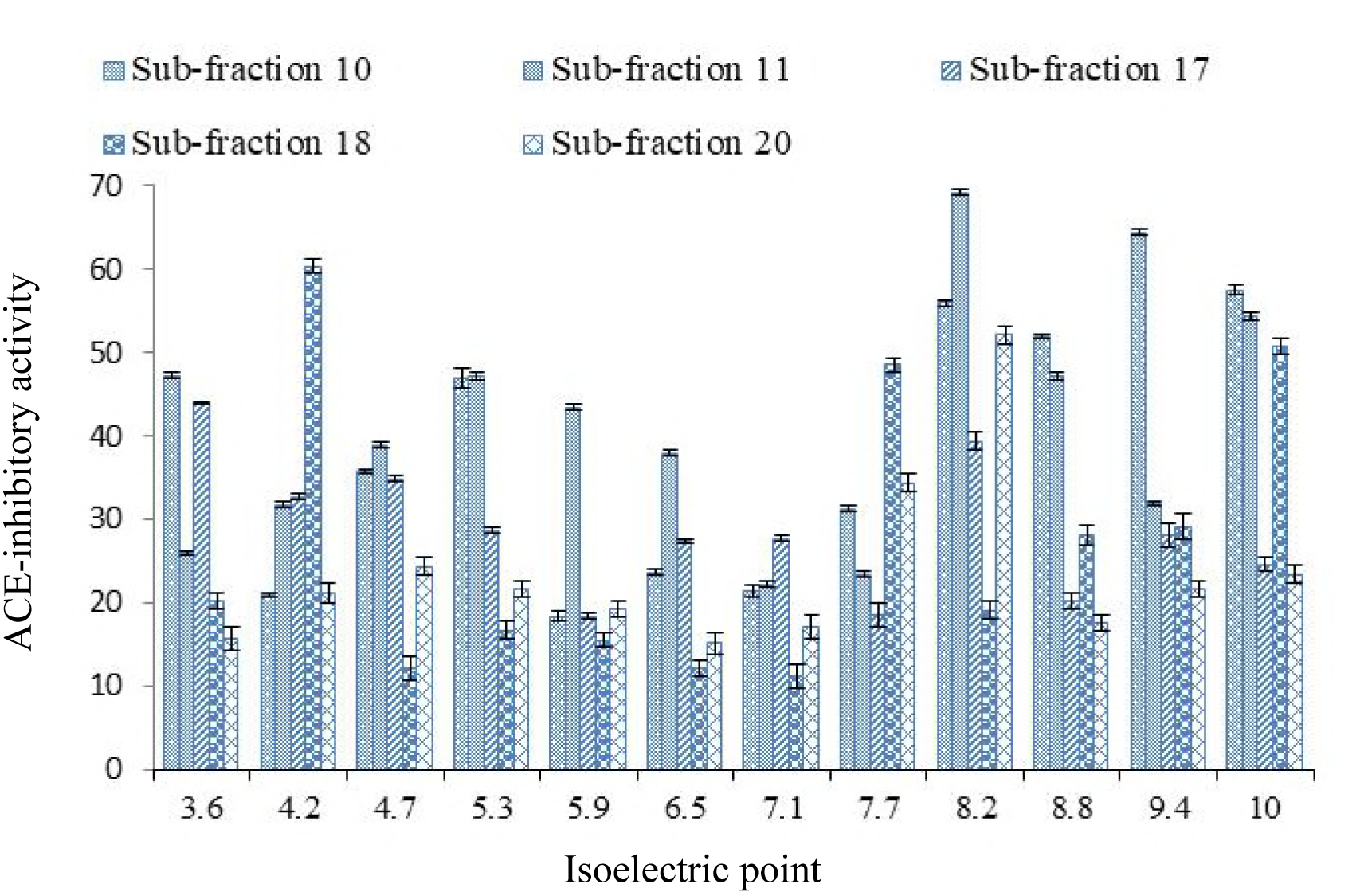
ACE-inhibitory activity of bromelain-generated stone fish protein hydrolysate fractions at different isoelectric points along a pH gradient (3-10).

Lastly, the IEF sub-fractions 11, 9, 1, 2 and 9 from the RP-HPLC fractions 10, 11, 17, 18 and 20 showed the highest ACE-inhibitory activity of 64.49%, 69.21%, 43.92%, 60.43% and 52.11%, respectively (Table 1) and were selected for peptide sequencing based on QTOF-LC/MS.

The selected IEF sub-fractions were subjected to Q-TOF LC/MS for the sequencing and identification of the potent peptides based on amino acids composition. The process involved a system of U-HPLC coupled to a Q-TOF mass spectrometer to separate and identify the individual peptide sequences present.

The results for the MS/MS spectra, ion tables, MS/MS fragments and standard error of the potent peptide sequences with mw < 1000 Da are shown in Figure 3.

**Figure 3.**
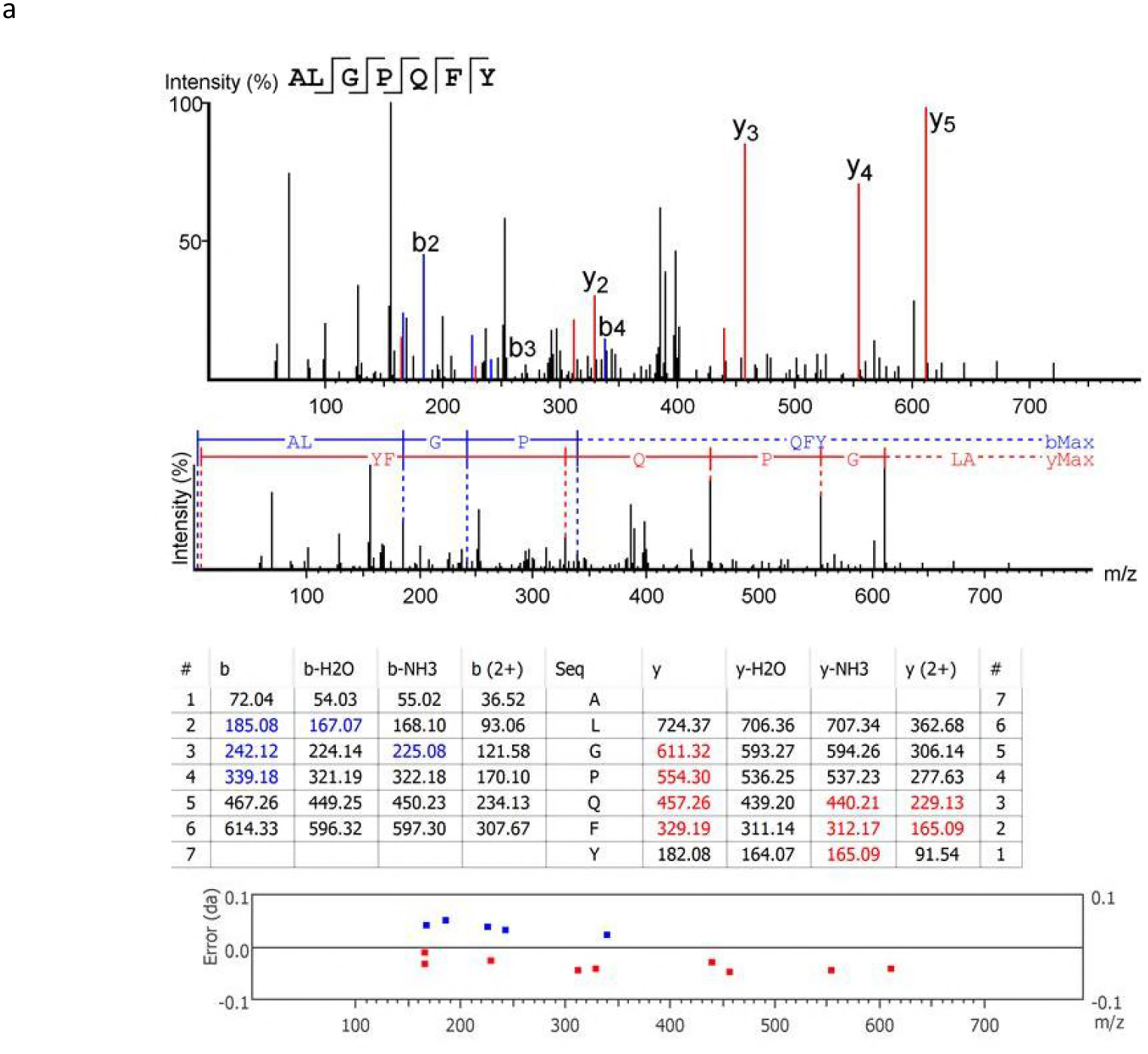
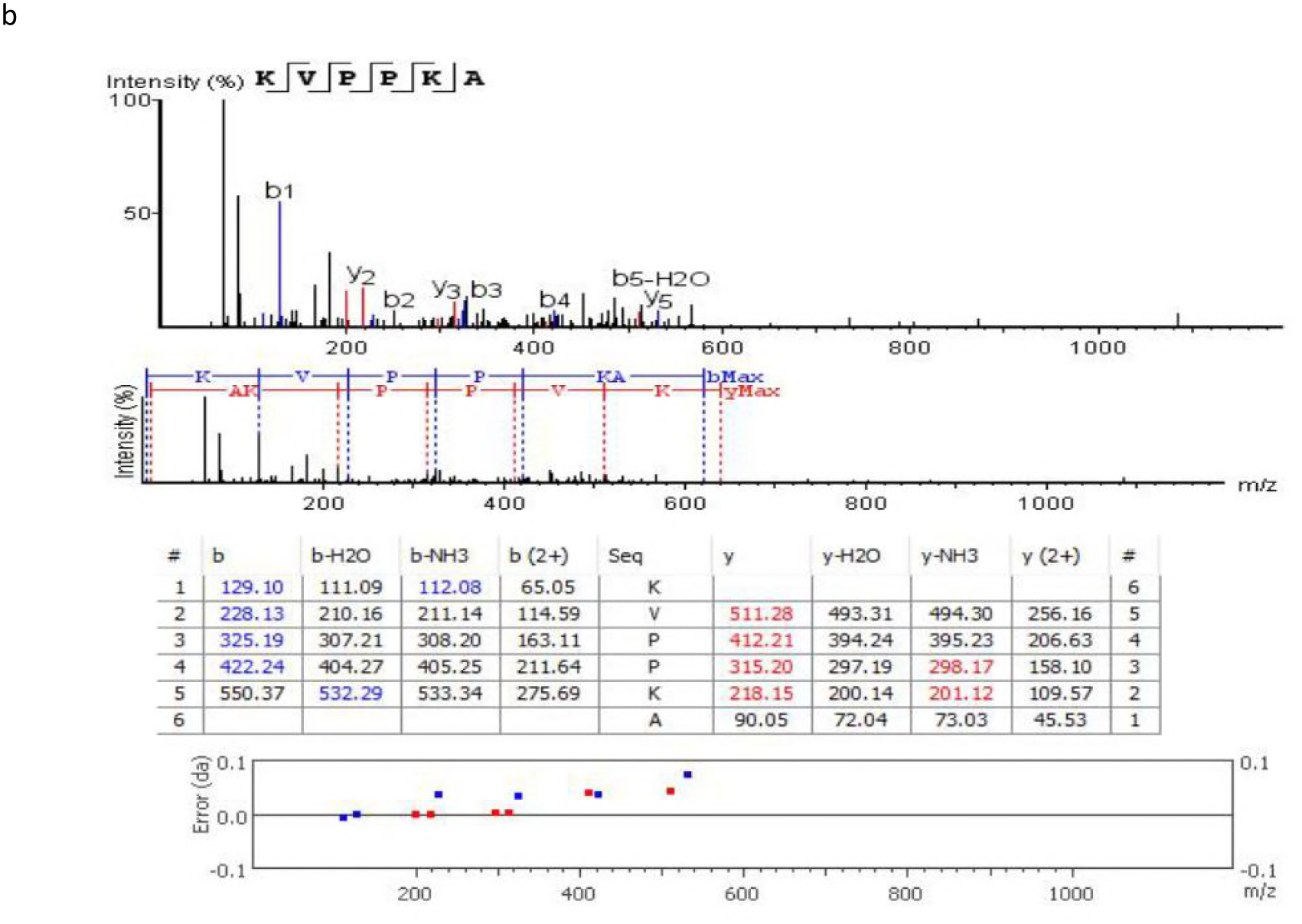
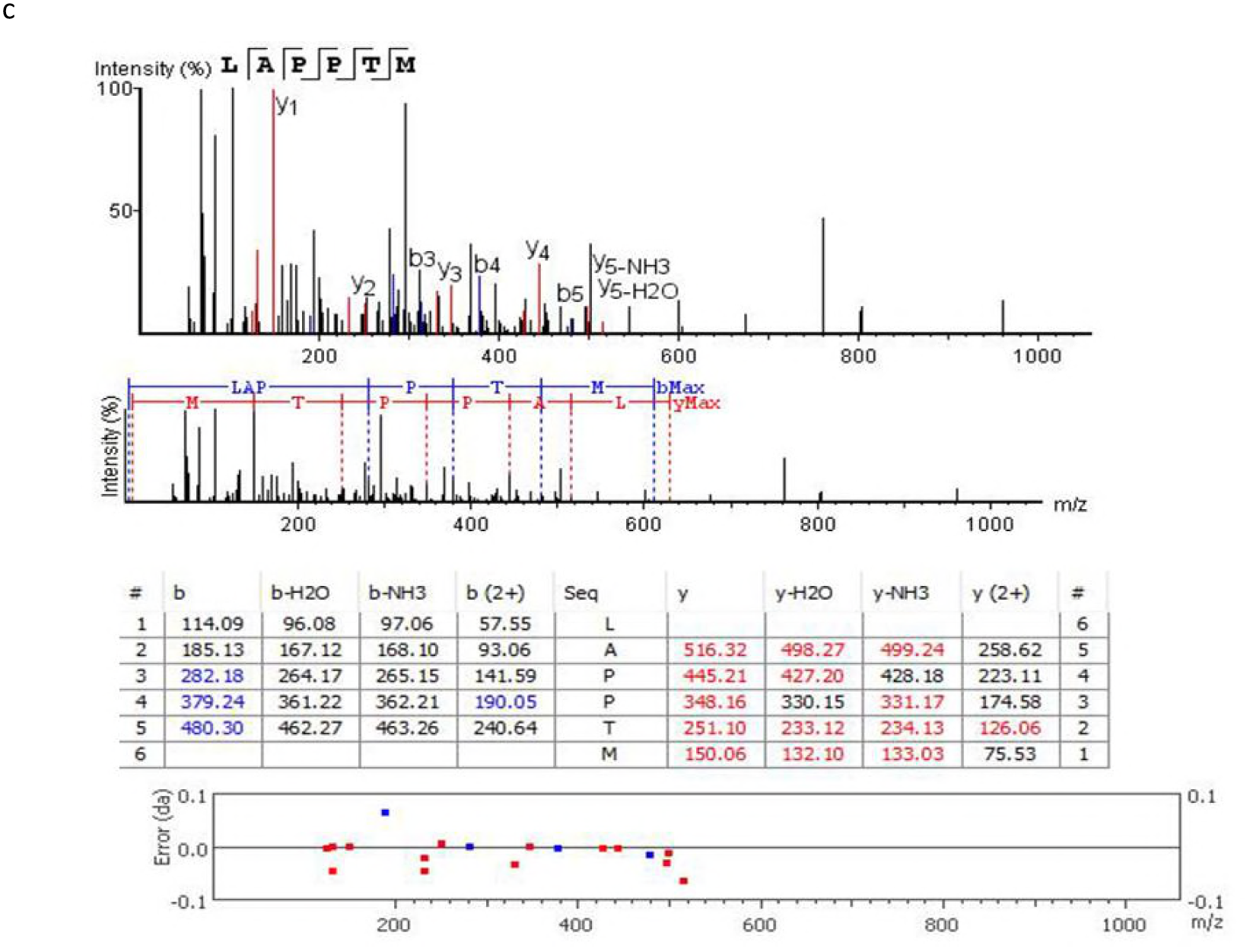
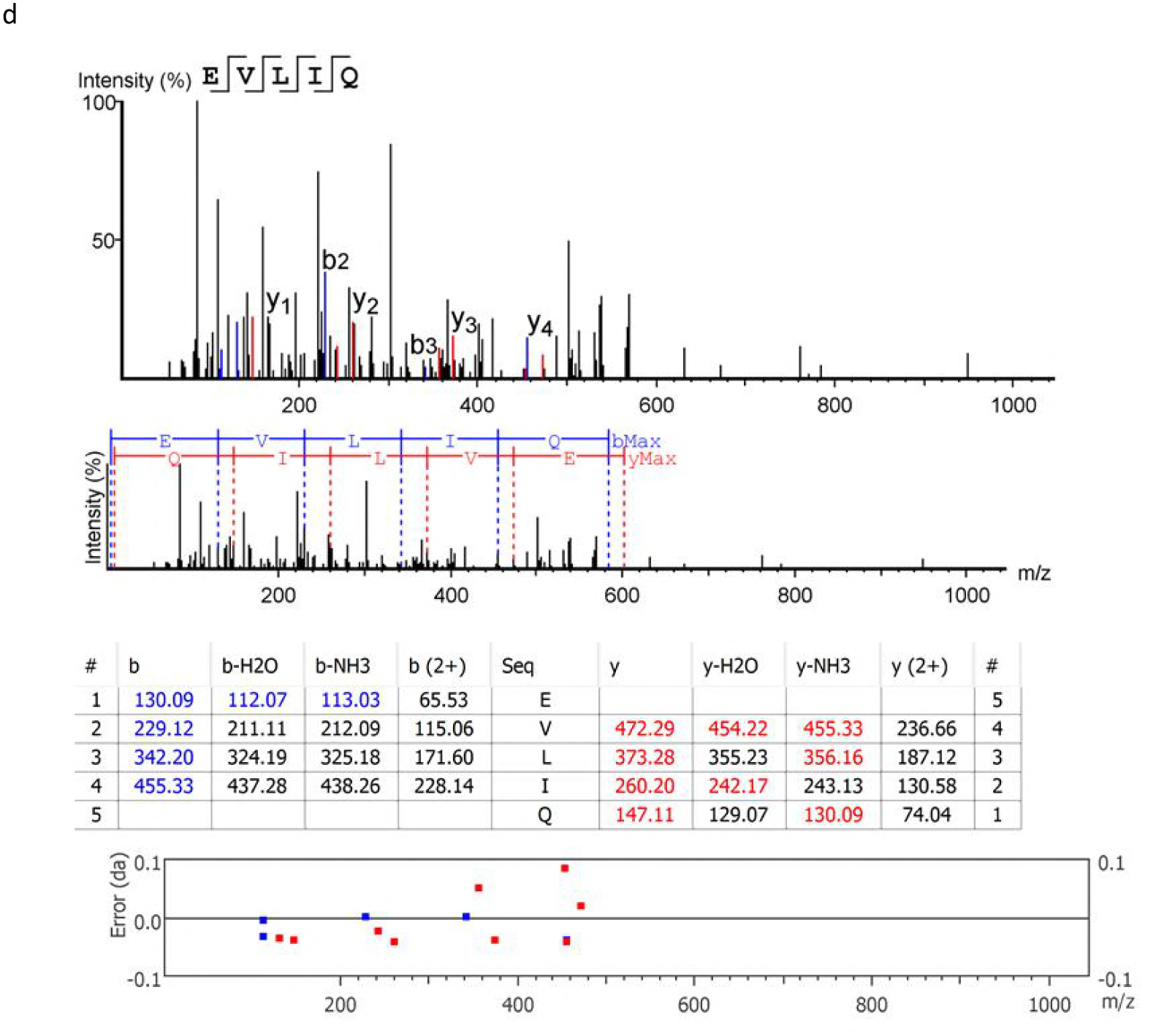
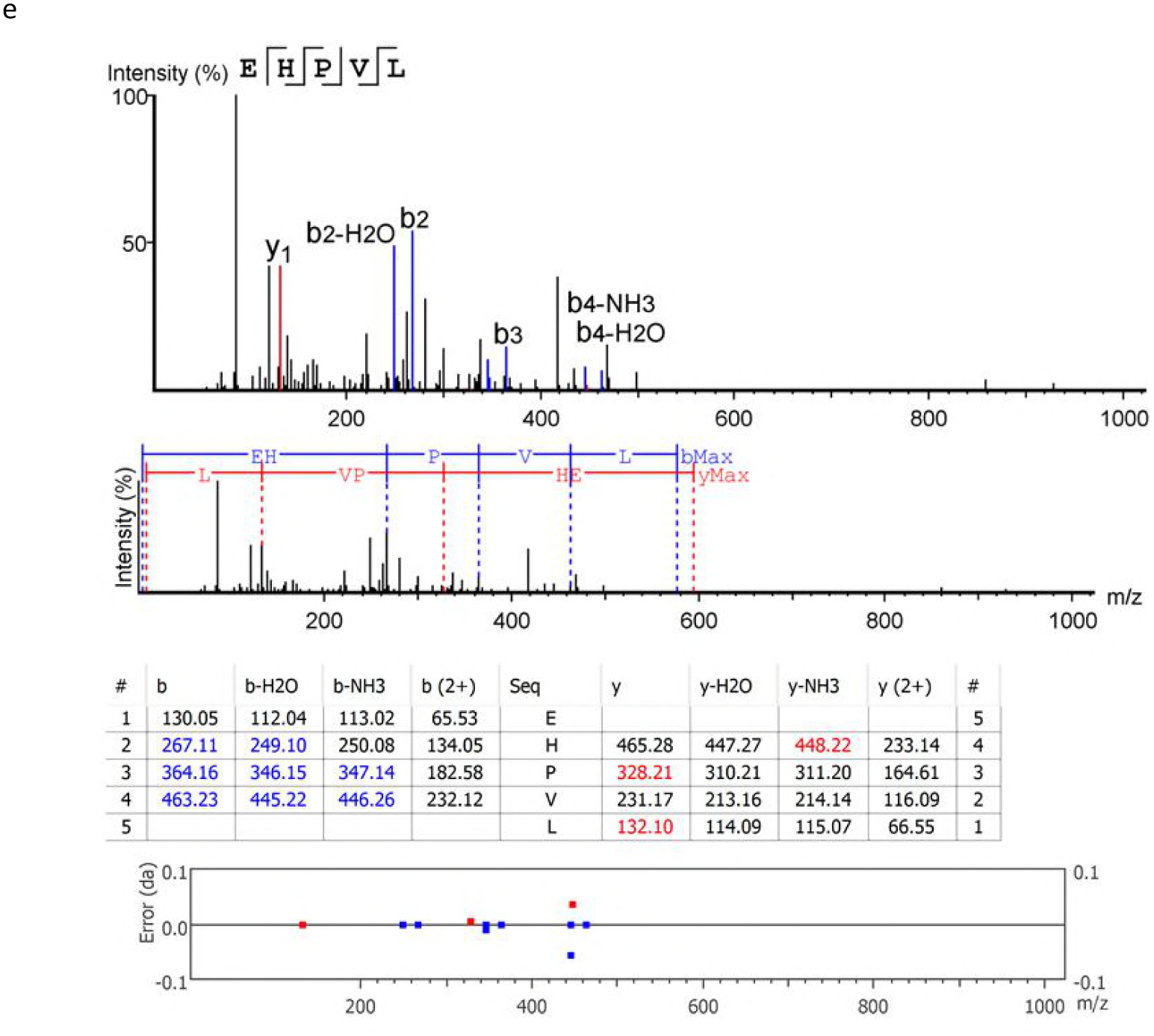
MS/MS spectra, ion tables, MS/MS fragments and standard error of the potent peptide sequences with mw < 1000 Da.

The identification, peptide sequence molecular formula confirmation and molecular ions mass accuracy determination were performed at a high resolution of 60,000. The MS/MS fragments and the ion tables have also been elucidated (Figure 3). The fragmentation involved ion peaks with two or more charges. High quality MS/MS spectra obtained for the peptide fractions were used for the identification of the peptide sequences.

As shown in Table 2, a total of five potent ACE-inhibitory peptide sequences with 5 to 7 amino acids residues and molecular weight < 1000 Da including ALGPQFY (794.44 Da), KVPPKA (638.88 Da), LAPPTM (628.85 Da), EVLIQ (600.77 Da) and EHPVL (593.74 Da) were identified by LC-MS and were found to be derivative of fragment spectra of high signal-to-noise ratio with complete or near-complete backbone fragmentation and low data error (≤ 0.1 da).

**Table 2.** Characteristics of the Potent ACE-inhibitory peptide sequences identified from the RP-HPLC fractions and IEF sub-fractions

| Peptide sequence | ACE-inhibitory activity at 10 mM (%) | IC<sub>50</sub> (mM) | Average mass (Da) | Number of amino acid residues | % Hydrophobicity/Hydrophilicity ratio of the constituent amino acids |  |  |  |
| --- | --- | --- | --- | --- | --- | --- | --- | --- |
|  |  |  |  |  | Acidic | Basic | Hydrophobic | Neutral |
| ALGPQFY | 94.43±2.053<sup>a</sup> | 0.012±0.001<sup>e</sup> | 794.44 | 7 | 0 | 0 | 57.14 | 42.86 |
| KVPPKA | 81.38±1.776<sup>b</sup> | 0.980±0.004<sup>d</sup> | 638.88 | 6 | 0 | 33.33 | 66.67 | 0 |
| LAPPTM | 72.39±1.993<sup>c</sup> | 1.31±0.017<sup>c</sup> | 628.85 | 6 | 0 | 0 | 83.33 | 16.67 |
| EVLIQ | 62.11±2.146<sup>d</sup> | 1.44±0.017<sup>b</sup> | 600.77 | 5 | 20 | 0 | 60.00 | 20 |
| EHPVL | 54.80±2.168<sup>e</sup> | 1.68±0.046<sup>a</sup> | 593.74 | 5 | 20 | 20 | 60.00 | 0 |

Results are expressed as mean ± SD of triplicate observations. Mean values of ACE-inhibitory activities and IC<sub>50</sub> of the peptides within the same column which carry different superscript letters are significantly different at $P \leq 0.05$ .

The peptides ALGPQFY, LAPPTM, EVLIQ and EHPVL have their PI in the acidic pH (3.6 – 6.5) while only peptide KVPPKA has its PI at the basic pH 10. All the peptide sequences have not been previously reported and are considered to be novel. The percentage hydrophobicity/hydrophilicity ratios of the peptides which affect their interaction with and subsequent effect on the active site of ACE have been indicated. The peptides were synthesized and their IC_50_ values were determined (Table 2).

Results are expressed as mean ± SD of triplicate observations. Mean values of ACE-inhibitory activities and IC_50_ of the peptides within the same column which carry different superscript letters are significantly different at P ≤ 0.05.

As previously reported, ACE-inhibitory activity of peptides is affected by their amino acids content. Potent ACE-inhibitory peptides contain short amino acids sequence between 2-12 residues and crystallographic studies have shown that large peptides may not bind to the active sites of ACE (23 - 25). The type of amino acids could be more relevant than the length of the peptide sequence. In this regard, highly acidic amino acids such as Asp and Glu may lead to a peptide with a net negative charge whose interaction with ACE could chelate zinc atoms in its active center to halt its activity (26). Presence of C-terminal hydrophobic tripeptides consisting of aromatic amino acids such as phe, tyr and tryp or N-terminal tripeptides containing branched chain aliphatic amino acids like val, leu and Ile resulted in high ACE-inhibitory activity and low IC_50_ of peptides (4,16, 27).

Moreover, highly hydrophilic peptides behave as weak ACE-inhibitors because they cannot properly fit to the active site of ACE. But, peptides with combined hydrophilic– hydrophobic property can favorably interact with the active center of ACE to inhibit its activity (16).

Majority of the previously reported ACE-inhibitory peptides derived from food sources contain pro and aromatic amino acids such as Tyr, Tryp or Phe at their C terminal (4, 16, 24, 28) and/or branched aliphatic amino acids such as Val, Leu, Ile at their N terminal (23). The positively charged amino acids arginine (R) and lysine (K) present at the C-terminal were also found to contribute to the ACE inhibitory potency of peptides (29). Thus, the observed differences in the ACE-inhibitory capacity exhibited by peptides can be attributed to the amino acids composition of their C and N terminal tripeptides (8).

According to our finding, the five peptides contained QFY, PKA, PTM, LIQ and PVL as their C-terminal tripeptide sequences with IC_50_ values of 0.012 mM, 0.980 mM, 1.31 mM, 1.44 mM and 1.68 mM, respectively. The peptides KVPPKA, LAPPTM and EHPVL contained Pro in the third position of their C-terminal tripeptides which might have contributed to their observed ACE-inhibitory effects. The additional Pro residue at the fourth positions of KVPPKA and LAPPTM could be responsible for their higher potency compared to EHPVL. Similarly, the presence of pro at the C-terminal has been reported to contribute to the ACE-inhibitory activity of peptides (30, 31). Moreover, the presence of lysine in the second position of KVPPKA could have strengthened its ACE-inhibitory effect. As previously reported, the ACE-inhibitory effect of peptide have been related to C-terminal arginine or lysine (32 - 35). Hence, in addition to terminal tripeptide sequence, the ACE-inhibitory capacity of peptides could be affected by the type of amino acid present at any other position in the sequence. Furthermore, ACE consists of two domains, which are N-domain and C-domain and they each contain a zinc cofactor binding active site (36). Therefore, ACE can be inhibited by metal chelating agents (26). The peptides ALGPQFY and EVLIQ share Glut residue at position 3 and 1 of their C-terminal sequences, respectively. Glutamic acid at the C-terminal of peptide contributed to its ACE-inhibitory potency for its tendency to chelate zinc in the active centre of ACE (37, 38). Consequently, the higher ACE-inhibitory activity observed for ALGPQFY could be ascribed to the presence of C-terminal aromatic amino acid residues (Phe and Tyr) and proline at the fourth position. Therefore, the ACE-inhibitory effect due to EVLIQ will largely relate to the presence of the branched chain aliphatic amino acid residues (Val, Leu and Ile) in its second, third and fourth N-terminal positions, respectively. Accordingly, the presence of Ala and/or Val in the N-terminal sequences of ALGPQFY, LAPPTM, KVPPKA and EVLIQ might have also contributed to their ACE-inhibitory potencies. This can be supported by the previous observation where branched-chain aliphatic amino acids present at the N-terminal were reported to affect competitive binding of peptide to the active site of ACE (39, 40). Thus, a peptide with branched aliphatic Ala, Val, Leu and Ile residues at its N-terminus, a basic arginine residue in the middle, and aromatic phe, tyr and tryp residues at its C-terminus could exert higher inhibitory activity against ACE (41).

In a related finding, bromelain-generated peptide sequence Ala-His-Leu-Leu with high ACE-inhibitory effect (IC_50_ of 18.2 ± 0.9 μg/mL) has been reported from *Misgurnus anguillicaudatus* tissue protein (42) and the IC_50_ value was found to be similar to the one obtained in the present study.

Thus, the ACE-inhibitory effect of peptides depend on the presence of hydrophobic and aromatic amino acid residues at one or more C-terminal position of peptides and/or N terminal branched aliphatic amino acids side-chains residues (35, 43). Potent ACE-inhibitory peptides were mostly characterized by the presence of Phe, Tyr, and Tryp and Pro at one or more C-terminal positions (35, 44). Furthermore, presence of Proline and hydroxyl proline was found to improve peptide resistance against gastrointestinal digestion and maintained higher bioavailability (45).

Even though the structure-activity relationship has not been well established for ACE-inhibitory peptides, it can be reliably stated that the inhibitory capacity of the peptides is greatly influenced by composition and orientation of amino acids within their sequences.

### *In silico* molecular docking of the inhibitory peptides and ACE

In the present study, a GlideXP software (Schrödinger, LLC) was use to dock ACE-inhibitory peptides as small molecular structures (ligands) to the ACE as a large molecular structure (protein) so as to deduce the optimum poses which favors the formation of stable interaction between them based on a scoring technique. The tendency of a particular peptide (ligand) to bind the protein receptor (ACE) at a specific conformation is rated by the GlideXP according to a rigid receptor approximation approach. Accordingly, the ligand-protein (peptide-ACE) fitting is governed by the potential to form hydrogen bonds, electrostatic interactions and to attain an acceptable root mean square deviation in comparison to the native complex (46, 48).

The GlideScore referred to an empirical scoring function that has been optimized for docking accuracy and binding affinity prediction. A more negative score signifies a stronger binding and a more stable ligand-protein (peptide-ACE) interaction (49).

Table 3 shows the estimated GlideScores and binding energies for the five stone fish-derived ACE-inhibitory peptides and that of the synthetic ACE-inhibitor (captopril).

**Table 3.** Estimated values of the GlideScore and binding energy for the best poses determined by molecular docking of stonefish-derived ACE-inhibitory peptides to the binding site of ACE.

| Ligands (Peptides) | GlideScore (kJ/mol) | Binding energy of ligand to receptor (kJ/mol) | ACE amino acids residues in coordination with $Zn^{2+}$ |
| --- | --- | --- | --- |
| ALGPQFY | -10.374 | -62.089 | Glu, His, Tyr |
| KVPPKA | -10.132 | -61.953 | His, Tyr |
| LAPPTM | -9.154 | -54.564 | His, Tyr |
| EVLIQ | -8.736 | -47.902 | Arg, Asp, His |
| EHPVL | -8.635 | -44.389 | Glut, His, Tyr |
| Captopril | -3.182 | -34.371 | Glut, His |

A good correlation was observed between the calculated ACE-inhibitory effect and the Glide score of the peptides. Peptide ALGPQFY with least IC_50_ value and stronger inhibitory effect on ACE showed the highest magnitude of negative GlideSscore of −10.374 kj/mol. Similarly, peptide EHPVL with highest IC_50_ value and lower inhibitory effect on ACE demonstrated the least negative GlideScore of −8.635 kj/mol.

The observed variation in the Glide score and ACE-inhibitory effect might be related to the stability of the peptide-ACE complex structure as affected by hydrogen bonds and electrostatic interactions with amino acids and zinc-cofactor in the ACE binding domain.

As shown in Table 3, the lower the binding energy the greater the stability of the peptide-ACE complex and the higher the inhibitory effect of the peptides on ACE. Thus, peptide ALGPQFY with the lowest binding energy of −62.089 kj/mol had the highest ACE-inhibitory (IC_50_, 0.012mM) effect whereas the peptide EHPVL with the highest binding energy of −44.389 kj/mol had the least ACE-inhibitory effect (IC_50_, 1.68 mM).

The atomic interactions between the amino acids residues of the peptides and that from the ACE within a distance of 3.5 Å also contributed to the stability of the peptide-ACE complex (50, 51). Right space orientation results in strong bond formation with the peptide in close proximity to ACE. The distance between the C-terminal carbonyl oxygen on peptide and Zn^2+^ cofactor on ACE binding domain affect the rate of peptide inhibition on the ACE (52). The shorter the distance the stronger the interaction and the higher the inhibitory effect (53). Thus, peptide ALGPQFY might have exhibited its lowest IC_50_ value and highest magnitude of the negative GlideScore via shorter C-terminal carbonyl oxygen - Zn^2+^ interaction and stronger atomic interaction between its amino acids side chain residues and those on ACE within a 3.5 Å distance.

The 2D structures presented in Figure 4 illustrate the poses for the binding of each of the five peptides and captopril (reference drug) to ACE. The different types of bonds formed, the amino acids and Zinc-cofactor in the ACE binding domain which are involved in its interaction with the peptides have also been indicated.

**Figure 4.**
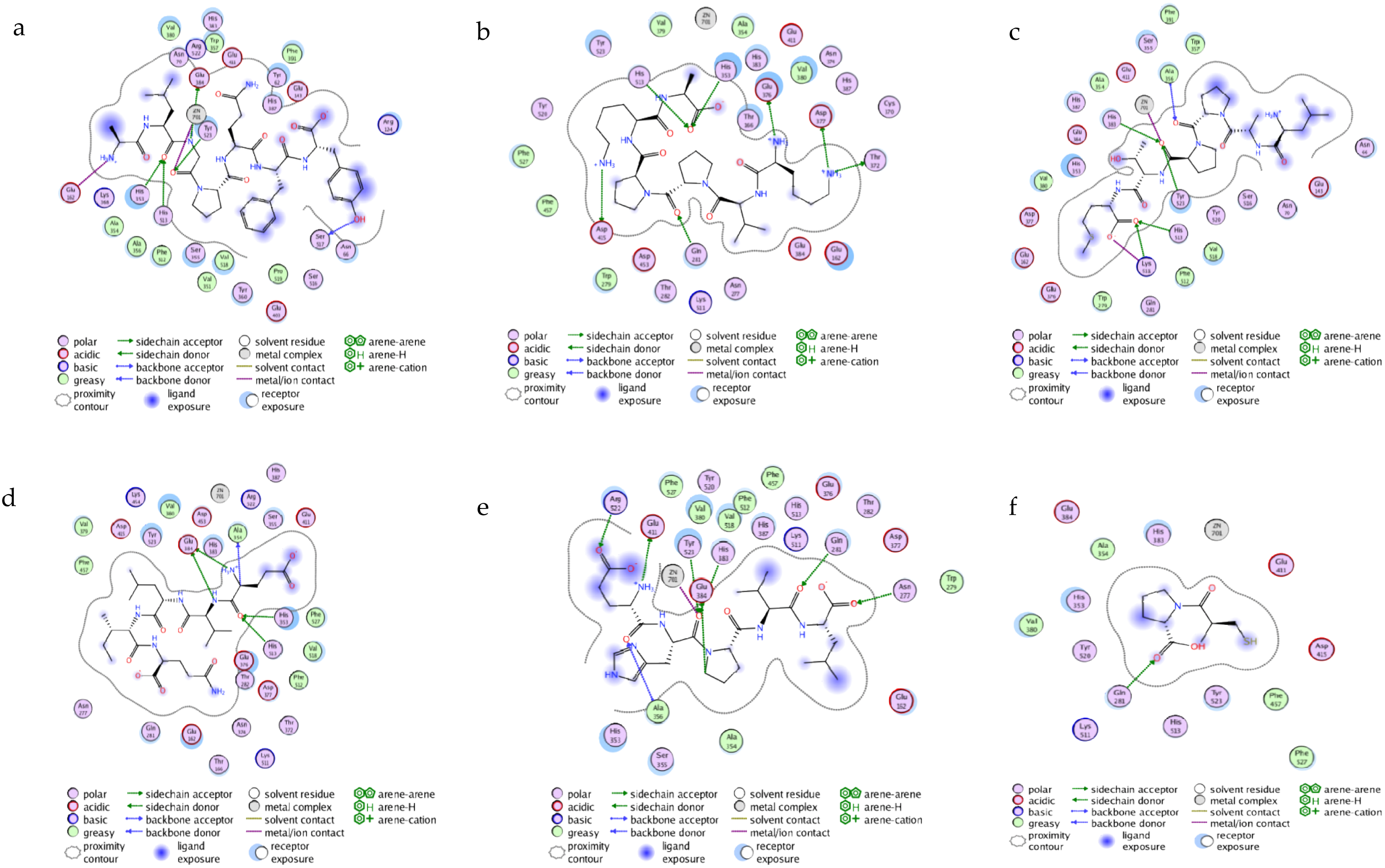
Predicted binding site for the 2D interaction of stone fish-derived ACE-inhibitory peptides with molecular surface of ACE; (a) ALGPQFY; (b) KVPPKA; (c) LAPPTM; (d) EVLIQ; (e) EHPVL and (f) Captopril as predicted by Schrödinger software

In Figure 4 (f), the captopril in the captopril-ACE complex is stabilized through hydrogen bond formed by its carbonyl oxygen with amino acids residue (Gln 281) on ACE and electrostatic linkage formed between the negatively charge thiol group in captopril with Zn^2+^ on the ACE coordinated through Glu 411 and via π-bonding between the nitrogen group on pyrrolidine ring of captopril and phenyl ring of Tyr 523 on ACE (54). Similarly, the peptide-ACE (ligand-protein) interaction is partly stabilized via multiple hydrogen bonds interaction with ACE amino acids residues and partly enhanced by electrostatic interactions between the amino acid side chain residues and/or Zinc-cofactor on the ACE binding site and the amino acids groups of the peptides. However, the type of ACE amino acids involved in hydrogen bond with those residues in the peptides could have greatly influenced the stability of the peptide-ACE complex and the inhibitory effect of the peptide on ACE. Here, the most active peptides (ALGPQFY and KVPPKA) formed hydrogen bonds with His353 and His513 amino acids which are critical residues to the activity of ACE (Table 3). Moreover, the higher inhibitory potency of ALGPQFY could be ascribed to the strong hydrogen bond formed between its aromatic amino acid Tyr 523 and the ACE binding center.

The free energy values obtained for the five peptides (−34.371 to −62.089 kj/mol) were found to be lower compared to the previously reported values of −13.97 to −21.59 kj/mol for peptides purified from Pacific cod skin gelatin-hydrolysates (55) and −31.46 to −39.91 kcal/mol for rape seed protein-generated peptides (51). These observations indicate higher affinity of the peptides identified in the present study for binding with ACE. However, comparably lower binding energy values of −92.91 to −114.06 kj/mol have been reported for hemp seed protein-generated peptides (50). We attributed the higher binding energy scores obtained for the peptides in this study to the effect of rigid-receptor approach used for calculation by the Glide program which deter good score for peptides with significant steric clashes in relation to its specified receptor conformation. However, in real situation the peptides may effectively bind with alternative conformation of the same ACE receptor protein (46, 48).

## Conclusion

In the present study, five novel ACE-inhibitory peptides with short amino acids sequences and mwt < 1000 Da were successfully identified from stone fish hydrolysates. The peptides including ALGPQFY (794.44 Da), KVPPKA (638.88 Da), LAPPTM (628.85 Da), EVLIQ (600.77 Da) and EHPVL (593.74 Da) exhibited strong inhibitory effect on ACE with IC_50_ values of 0.012 mM, 0.980 mM, 1.31 mM, 1.41 mM and 1.68 mM, respectively. The observed differences in the ACE-inhibitory effects of the peptides was found to be affected by their C and N terminal tripeptide sequences based on structure-activity relationship. Moreover, molecular docking studies showed that the peptide-ACE docking complex is stabilized through hydrogen bonds and electrostatic interactions between the amino acids side chain residues and/or zinc-cofactor in the ACE binding sites to the amino acid residues of the peptides. Thus, the results of the present study indicate the potential of stone fish protein hydrolysates as a suitable raw material for the industrial production of ACE-inhibitory peptides which could be used in effective treatment of hypertension.

## Acknowledgments

The authors acknowledged funding support by the Malaysian Ministry of Science, Technology and Innovation (MOSTI) under the project (No. 02-01-04-SF 2309).

